# Infrared videography of a subcutaneous knee tattoo as a simple and inexpensive method to overcome skin motion artifact in rodent kinematics

**DOI:** 10.1101/2022.08.22.504813

**Authors:** George Moukarzel, Bradley C. Rauscher, Chetan A. Patil, Andrew J. Spence

## Abstract

Kinematic analyses of rodent behavior are frequently used in neuroscience research, and commonly in spinal cord injury (SCI) studies. Unfortunately, skin motion artifact introduces significant errors into these data, because the skin is only loosely coupled to the underlying skeleton by connective tissue. In rats, these errors can be as large as 50-75%,as quantified by past work using x-ray fluoroscopy. Here we show that infrared videography of a subcutaneous tattoo can overcome skin motion artifact in rodent kinematics. The method yields data similar to gold standard x-ray fluoroscopy systems at a fraction of the cost, does not affect the animals’ locomotion, and results in markers that persist for at least 10 weeks. We found that, compared to a gold-standard x-ray fluoroscopy study that directly tracked the skeleton, our method reduced the error in mean hip angle from 17 ± 6.0 to 3.1 ± 2.4 degrees (mean ± SEM), and the root-mean-square (RMS) error across the mean hip angle waveform from 20 to 5.3 degrees (n=4 rats). The knee joint angle waveform derived from infra-red imaging tightly matched the shape of the x-ray waveform after allowing for a constant offset, having RMS error reduced from 8.1 to 1.2 degrees. The method stands to significantly reduce between-animal errors, and hence between laboratory errors, in these ubiquitous model systems, especially important in SCI studies where individuals are assigned to different treatments.

## Introduction

Kinematic analyses of biological movement give important insight into health and disease, especially in spinal cord injury (SCI) studies. Unfortunately, and in rodents particularly, the skin is only loosely attached to the skeleton, and thus estimating skeletal posture from markers drawn on or attached to the skin is prone to significant error, known as skin motion artifact (1–4). For rats, a prominent SCI model (5), this problem is especially acute at the knee joint, where hip and knee angles can have error rates of 50-75% (3, 6).

One method to correct these errors is triangulation (7). This uses the positions of other markers and the known, fixed lengths of bone segments to deduce where a joint should be. Triangulation can suffer from uniqueness problems, the requirement for nearby good markers, and the need for bone lengths, preferably individualized, however. Incorrect lengths can cause erroneous joint positions (6), even increasing joint location errors (2, 4). Fluoroscopy, or direct videography of the bones *in vivo* using x-rays, is currently the gold standard for measuring true joint locations (6, 8). These systems are expensive, however, on the order of $1M for a turnkey, biplanar system, and typically have limited capture volumes and require shielding and limited exposure times.

Here we present a simple, inexpensive technique to overcome the main source of hindlimb skin motion artifact in rats, by directly imaging the knee joint through the skin using infra-red videography of a subcutaneous tattooing.

## Methods

### Study design

All animal procedures were approved by the Temple University Institutional Animal Care and Use Committee under Animal Care and Usage Protocol #5003. Adult female Long-Evans rats were used. We define two types of marker in this study: skin-derived (SD) markers drawn on the skin using permanent marker; and tattoo-derived (TD) markers tattooed subcutaneously to the knee joint.

Baseline treadmill locomotion videos with skin markings were recorded for all rats before tattooing. These were compared with data gathered two weeks after the procedure to ensure that 1) the dark spot seen with infrared imaging was the tattoo and not shadow or other imaging artifact (Fig. 1D and E; Supplemental Video S2), and 2) that tattooing had not affected kinematics (Fig. 2G, H, I). To ensure that the tattooed marker is durable for the length of a typical SCI study, data were captured at 10 weeks following the procedure, and the ability of our DeepLabCut (9) network to track the marker with identical settings of itself and the apparatus was tested (Fig. 1F). Finally, to assess whether our method corrects for skin motion artifact, skin-derived and tattoo-derived kinematics from the same videos were analyzed and compared with gold standard results from an x-ray system (Fig. 2A, B, and C; (6)) as well as prior studies of rat kinematics in the literature (Supplemental Information Figure S1; (6, 10, 11)).

**Figure 1.**
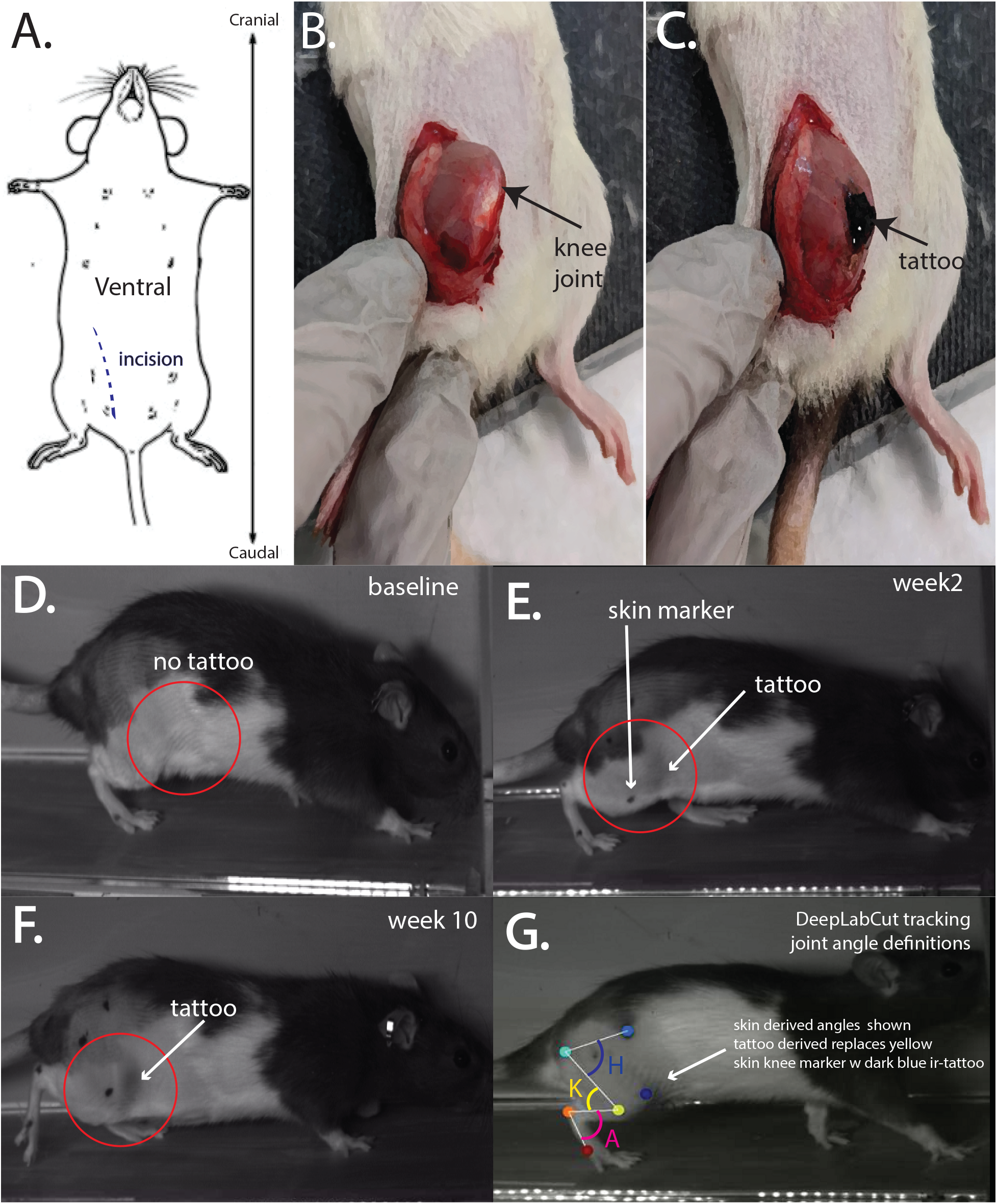
Subcutaneous tattooing of the knee and subsequent infrared videography gives more accurate tracking of the knee than skin markers, is durable, and reveals the extent of skin motion artifact. **A**. A 2-3 cm incision is made running over the inner thigh starting 5 mm above the knee and running medial to the knee joint to a point 2-3 mm below the knee. This incision location ensures that no scar tissue will form in the camera’s view. **B**. The knee is gently exposed through the incision, and small amounts of connective tissue are cleared. **C**. A tattooed knee joint. **D**. An infrared video of a rat prior to the tattooing procedure shows no marker on the knee joint. This shows that the dark region seen after tattooing is not shadow or other imaging artifact. **E**. The subcutaneous tattoo is seen in infrared. As the skin marker is also visible, infrared videography reveals the extent of skin motion artifact in this frame. The SD and TD knee markers do not coincide, and the knee joint has higher variability compared to the SD knee marker (See Supplemental Video S1). **F**. 10 weeks after the tattooing procedure was performed, the TD marker was still detected under the same settings of the experiment, demonstrating that the tattoo endures for the length of a typical SCI study. **G**. Illustration of the hip, knee and ankle angles derived from the tracked joint positions (H, K and A respectively). Markers were automatically tracked using DeepLabCut on the raw 2D videos, 3D marker positions were reconstructed, and then joint angles were computed.

**Figure 2.**
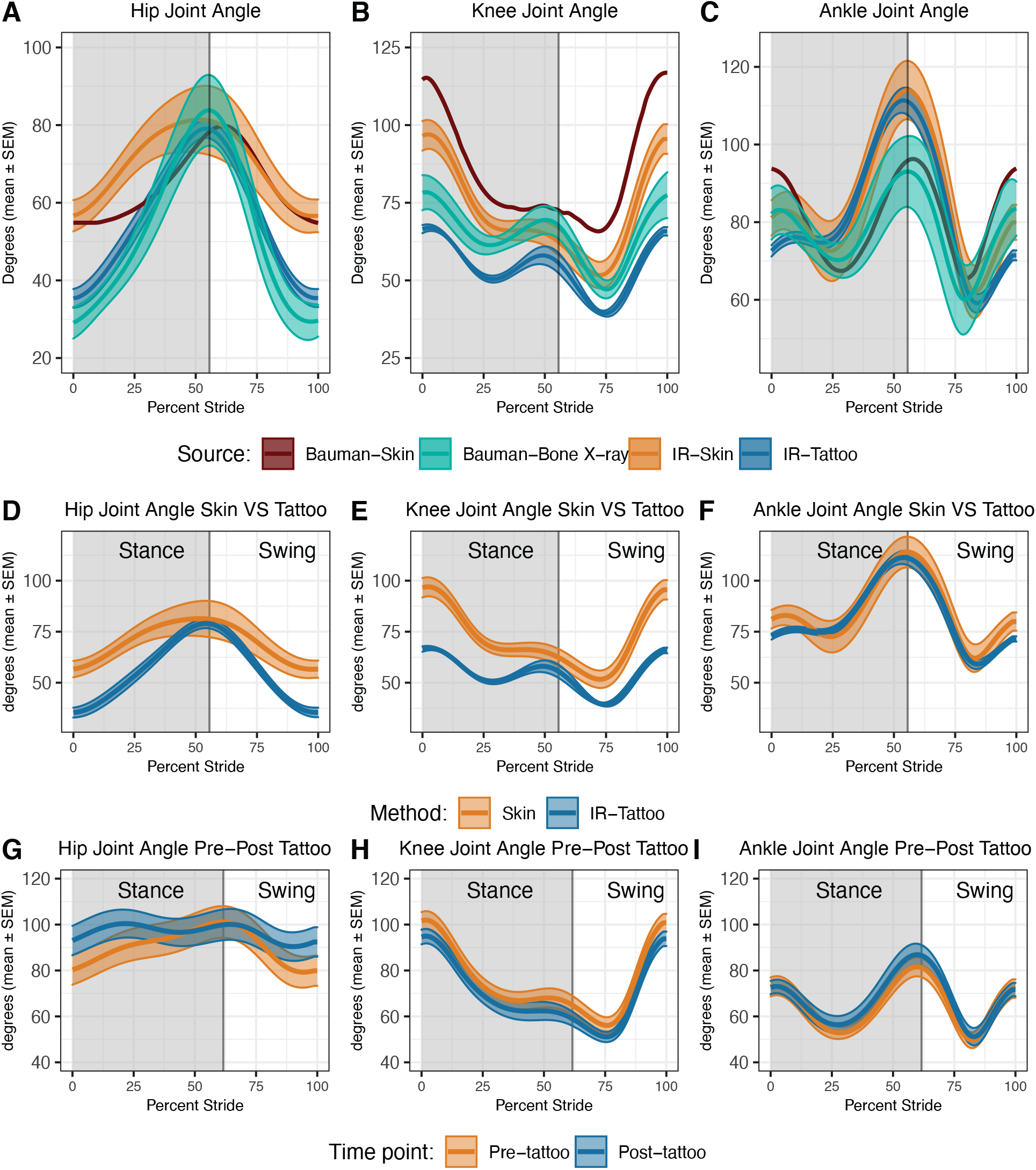
Infrared Imaging of the tattooed knee joint under the skin corrects for skin motion artifact when compared to x-ray fluoroscopy data (6) and does not affect the animals’ movement. **A**. Tattoo derived (TD) hip angles (IR-tattoo; dark blue) closely matched x-ray fluoroscopy data directly tracking the bones (Bauman-X-ray; teal) as compared to skin derived (SD) data from the same videos (IR-skin; tan). Skin derived data matched that in the Bauman study (Bauman-skin; dark brown). Mean hip angle difference was reduced from 17 ± 6.0 to 3.1 ± 2.4 degrees (IR-skin – Bauman-x-ray vs IR-tattoo – Bauman-X-ray; mean ± SEM; n=4 rats; median 40 strides per rat, min 15, max 48). Root-mean-square (RMS) error across the mean hip angle waveform was reduced from 20 to 5.3 degrees. **B**. The TD knee joint angle waveform tightly matched the shape of the x-ray waveform after allowing for a small constant offset, having RMS error reduced from 8.1 to 1.2 degrees **C**. TD ankle angle trended towards the x-ray data at liftoff but was not substantially altered, likely do to less dependence on the knee marker. **D, E, F**. Tattoo-derived kinematics revealed that a large range of motion of the knee joint under the skin causes the skin marker error. **D**. We found that the SD kinematics of the hip represented a 22.3 ± 1.64-degree overestimate of mean hip angle when compared to TD data, and that SD data underestimated the variability of the hip (standard deviation of the hip angle across percent stance within each stride increased by 28.3 ± 0.2). **E**. The SD data overestimated the knee angle at paw touchdown (0 and 100% stride) relative to TD data by 21.1± 3.2 degrees. **F**. As in past literature, we found TD ankle kinematics similar to SD kinematics, where the influence of knee skin-motion artifact is minimal, with only a 10.5 ± 4.3-degree difference at paw touchdown. The difference in knee marker position between TD and SD is highly noticeable throughout the stride, especially large during swing and early stance, and minimized at liftoff, for our skin marker position (Supplemental Videos S1 and S4). **G, H, I**. Tattooing the knee joint under the skin did not affect joint angle kinematics. Seven naïve female Long-Evans rats ran at three different speeds (16, 24 and 32 cm/s) before and two weeks after tattooing (32 cm/s data shown). Wilcoxon rank sum tests were carried out at each percent stance bin (median 36 strides per rat, min 22, max 61). No percent stance bins were significant at the p<0.05 level for any speed.

### Kinematic data capture from high speed infra-red video

All animals were placed under isoflurane anesthesia, shaved, palpated to locate landmarks, and marked with permanent marker over the anterior-superior iliac spine (ASIS), hip, knee, ankle, and toe joints, to create the SD markers (Fig. 1G). The animal was held as close as possible to mid-stance posture.

Three infrared (IR) light-emitting diode (LED) panels (CMVision; CM-IR130-198; ~$73) emitting at 850nm were used as light sources. Two high-speed IR cameras (Ximea XiQ MQ022RG-CM; ~$1800 each with lens; 250 frames/sec) were used to take synchronized videos. To control the treadmill speed, synchronize the cameras, and capture end-triggered video data, we employed a custom roboticized treadmill system previously developed (Supplementary Figure S2)(12–15). However, the method will work with any standard video based kinematics apparatus, assuming sufficient contrast of the IR tattoo.

The x-ray data presented in Bauman and Chang (6) and redrawn in Figure 2ABC are averages across a range of speeds, from 16.5–63.2 cm/s. We therefore present data at 48 cm/s, in the middle of this range, for comparison in these panels (Fig. 2ABC).

### Feature tracking and kinematic computation

Automatic marker tracking and reconstruction were carried out with DeepLabCut (9). Six features were tracked: the subcutaneous tattoo on the knee and the five SD markers (Fig 1G). Raw 2D marker coordinates were extracted from the two camera views and then reconstructed in 3D, followed by computation of joint angles, using a custom Python code-base (7). Cameras were calibrated with a custom LEGO calibration object and the free MATLAB TyDLT package (16). Because the available x-ray fluoroscopy data are 2D (6), we projected our data down to 2D by removing the z-coordinate. SD joint angles were computed using the coordinates of the permanent marker on the knee, whereas TD angles were computed using the coordinates of the knee tattoo. Joint angle data were first averaged across strides at each percent stance bin within each unique grouping of rat, imaging type, and speed, resulting in one average time series for each rat for each imaging type. The mean and standard error of these rat means was then computed separately for rats within each time point and/or imaging type (baseline or post-tattoo; skin-derived or tattoo-derived), as a function of percent stride bin, resulting in the plots in Figure 2, which are displayed as mean ± standard error. For comparisons of mean or standard deviations of joint angles, the mean joint angle was taken across percent stance of the final individual rat mean waveforms to produce one mean or standard deviation per rat per marker type, which was then compared to the same statistic taken across the x-ray waveform.

### Tattooing procedure

Tattoos were applied using an AIMS™ ATS-3 General Rodent Tattoo System. Animals were anesthetized during the procedure (isoflurane: 5% induction and 3% maintenance; 100% O2). A 2-3 cm incision running over the inner thigh starting 5 mm above the knee and running medial to the knee joint to a point 2-3 mm below the knee (Fig. 1ABC). A ventral incision ensured that scar tissue would not obstruct the marker during imaging. The right knee was exposed by sliding this incision up over the protruding knee joint. Small amounts of fascia and connective tissue were gently cut to expose the surface of the musculature directly attached to the knee. Black pigment #242 (AIMS Inc.) was tattooed on the knee area that covers the joint and connective tissue (Fig 1BC). The wound was closed with wound clips. Rats were given two weeks for recovery before running.

## Results

### Tattoo-derived kinematics correct for skin motion artifact when compared to gold-standard x-ray fluoroscopy data (Figure 2A, B, C; Supplemental Videos S1 and S4)

The difference in mean hip angle between our data (2 weeks post tattoo; 48 cm/s; Supplemental Videos S1 and S4) and x-ray fluoroscopy data (Fig. 2A) was reduced from 17 ± 6.0 to 3.1 ± 2.4 degrees (mean ± SEM; n=4 rats; median 40 strides per rat, min 15, max 48) by utilizing the infra-red visualized TD knee marker instead of the SD knee marker (in the same videos), and the root-mean-square (RMS) error across the mean hip angle waveform was reduced from 20 to 5.3 degrees (Fig 2A). The TD knee joint angle waveform tightly matched the shape of the x-ray waveform after allowing for a constant offset, having RMS error reduced from 8.1 to 1.2 degrees (Fig. 2B). TD ankle angle trended towards the x-ray data at liftoff but was not substantially different than SD (Fig 2C).

### Tattoo-derived kinematics reveal that a large range of motion of the knee joint under the skin drives skin marker error

The SD kinematics gave a 22.3 ± 1.64-degree overestimate of the mean hip angle when compared to TD (Fig. 2D). Further, the standard deviation of the hip angle across percent stance increased by 28.3 ± 0.2 degrees (Fig. 2D) in the TD data. On average, the hip angle standard deviation was 16.1 degrees when analyzing SD kinematics, compared to 44.5 degrees in TD kinematics. SD kinematics further gave a large overestimation of knee angle at paw touchdown (0 and 100% stride) of 21.1± 3.2 degrees (Fig. 2E). Similar to past literature, we found TD ankle kinematics resembled those of SD (Fig. 2F), where we observed only a 10.5 ±4.3-degree difference at touchdown. The difference in knee marker position was noticeable throughout the stride, especially large during swing and early stance, and minimized at liftoff.

### Tattooing the knee joint under the skin did not affect joint angle kinematics

Seven naïve female Long-Evans rats ran at three different speeds (16, 24 and 32 cm/s) before and two weeks after tattooing (Fig 2G, H, and I display 32 cm/s). All strides within each rat, speed, and time point were averaged to produce one waveform per joint angle per rat, and then Wilcoxon rank sum tests were carried out at each percent stance bin (median 36 strides per rat, min 22, max 61 for 32 cm/s). No percent stance bins were significant at the p<0.05 level for any speed.

### The subcutaneous knee tattoo persists long enough for longitudinal studies

Animals were recorded running at 16, 24 and 32 cm/s at both two weeks and 10 weeks following the tattoo with identical imaging settings (Fig. 1F; Supplemental Video S3). We analyzed the 10 week videos with the same DeepLabCut network used at two weeks without refinement. The TD knee joint was detected at p>0.95 in 89% ± 6% of the time at 10 weeks (8 rats, 9 trials, 2 cameras at 1000 frames per trial comprising 144,000 frames) as compared to 88 ± 12% at two weeks (6 rats, 9 trials, 2 cameras comprising 108,000 frames total). We note that this probability of detection is readily increased by improving the DLC network; this network was trained with two videos per rat, 50 frames per video, for only 500 frames total. A prior network applied to similar infrared data with 200 frames per rat totaling 4,800 training frames achieved 96.5% detection. The marker remained visible to inspection in the animals that remained willing to run (three animals) at 16 and 26 weeks post tattoo (data not shown).

## Discussion

We describe a novel method to more accurately track the knee joint in the moving rat without x-ray imaging or triangulation. We show that this method improves the accuracy of knee marker tracking and resulting joint angle data, is durable, and does not alter the kinematics of the animal. The main limitations of the method are that it requires a recovery surgery and infra-red cameras. The surgery is minimally invasive and short (10 min), however, and infra-red cameras are relatively inexpensive (~$1500 per camera). The data analysis utilizes standard video workflows.

Our results illustrate how between-animal or between-marker application errors result from placement of the knee marker on the skin, and large subcutaneous knee joint movement (Supplemental Videos S1 and S4). While every effort is made to repeatably mark the animal, inevitable differences in skin marker location will cause DC-shifts, and potentially other nonlinear changes. Comparing SD kinematics from our study with three others in the literature shows these shifts (Supplementary Figure S2) (6, 10, 11). In addition, because the knee moves so substantially beneath the skin (Supplementary Video S1), no skin marker placement can be accurate throughout the stride, as indicated by the increased variability of the TD hip kinematics (e.g. Fig. 2A, D).

A potential reason for the remaining discrepancies of our knee and ankle TD joint angles compared with the x-ray data is that our knee marker may lie on the rostral surface of the knee rather than the lateral aspect, which could more accurately represent the joint center. This could be improved with better tattoo placement.

Future work could seek to improve contrast, through fluorescence and/or novel biomaterials, and should expand to more strains of rat, the forelimbs, mice, and other animal models where ethical. The tattoo could be better localized. Finally, estimation of the between-animal variability of the knee tattoo against x-ray fluoroscopy would be beneficial.

Because the method is inexpensive and scalable to many animals and long bouts of locomotion, we hope that it will aid in establishing large databases of shared kinematic data with less between-lab variability.

## Supporting information

Supplemental Video S1 Skin vs Tattoo

Supplemental Video S2 - No Tattoo Control

Supplemental Video S3 - 10 Week Longevity Test

Supplemental Video S4 - DeepLabCut Tracking

## Acknowledgments

This is work was supported by NIH Grant #1R01NS114007-01A1 to AJS. We thank NVIDIA corporation for their generous donation of the Titan Xp GPU used in this work. We thank the many undergraduate research assistants in the Spence Lab, the staff and administration of the Bioengineering Department, and members of the Smith and Lemay labs for their assistance.

## Author Contributions

AJS conceived of the approach, GM lead and managed all aspects of the study, gathered and analyzed data, and wrote the manuscript draft. CP suggested improvements to contrast and imaging; GM and AJS developed the surgical approach; BR carried out data collection and analysis, and further suggested refinements to the approach.

## Supplemental Data

Supplemental Video S1: Video of rat with tattoo showing discrepancy between skin and tattoo markers.

Supplemental Video S2: Video of rat at baseline before tattooing, with identical imaging parameters as Video S1, showing absence of tattoo, to confirm that the dark region is indeed the tattoo and not shadow or other imaging artifact.

Supplemental Video S3: Video of a rat at 10 weeks post tattoo demonstrating longevity of the tattoo.

Supplemental Video S4: Example of automated DeepLabCut tracking of skin and tattooed markers.

Supplemental Information

