## Supplementary Text and Figures for "Infrared videography of a subcutaneous knee tattoo as a simple and inexpensive method to overcome skin motion artifact in rodent kinematics"

### Supplemental Information

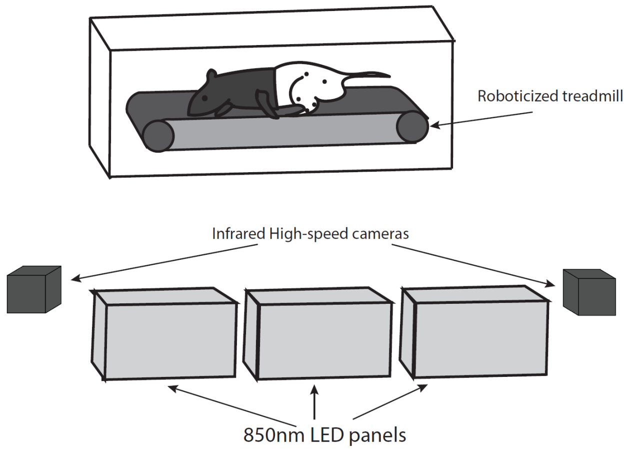


**Supplementary Figure S1.** Kinematics apparatus used in this study. Animals were marked with permanent marker over the anterior-superior iliac spine (ASIS), hip, knee, ankle, and toe joints, to create the skin-derived (SD) markers. Three infrared (IR) light-emitting diode (LED) panels (CMVision; CM-IR130-1983) emitting at 850nm were used as light sources. Two high-speed IR cameras (Ximea XiQ MQ022RG-CM; 250 frames/sec) were used to take synchronized videos. To control the treadmill speed, synchronize the cameras, and capture end-triggered video data, we employed a previously developed roboticized treadmill system (1–4).

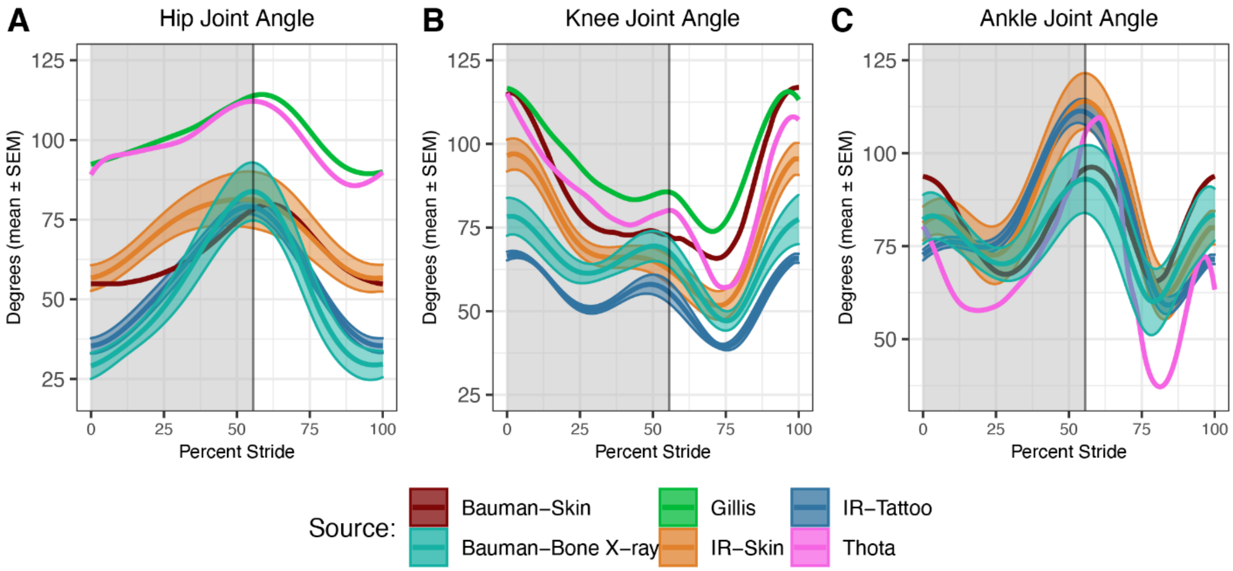

**Supplementary Figure S2.** Comparison of skin and tattoo data with additional studies from the literature (5–7). Figure is identical to Figure 2 (see caption), with the addition of redrawn data from studies by Gillis and Biewener, 2001, and Thota, 2004. The data from Bauman and Chang (2009; cf Fig 4ABC of that paper) are from six adult male Sprague-Dawley rats weighing 250 +- 33g running at speeds ranging from 16.5 to 63.2 cm/s, with skin markers placed under anesthesia in a manner similar to this study. The data from Bauman and Chang are two-dimensional, derived from a single camera view. The data from Gillis and Biewener (2001) are from ~40 cm/s walking gait locomotion in six female Sprague Dawley rats weighing 225-260g (cf Fig 2A,E of that paper). The data from Thota are at 30-35 cm/s in young adult (71 days) female Long-Evans rats weighting 211+-15 g, gathered in 3D using a Peak Motus system.
